# Frequent sexual reproduction limits adaptation in outcrossed populations of the alga *Chlamydomonas reinhardtii*

**DOI:** 10.1101/2022.07.01.498510

**Authors:** Felix Moerman, Nick Colegrave

## Abstract

Sexual reproduction can facilitate adaptation by reshuffling genetic variation. However, sexual reproduction can also bear costs. Such costs come in two forms: direct costs and evolutionary costs. Direct costs are associated with the cost of producing males (twofold cost of sex), the cost of meiosis, and the typically slower cell division during sexual reproduction of single-celled organisms. Evolutionary costs occur when too frequent sexual reproduction would hinder adaptation, by breaking apart adaptive allele combinations. Whereas the direct costs of sexual reproduction have been studied repeatedly in theoretical studies, the evolutionary costs of sex remain less well understood. We investigate here how the frequency of sexual reproduction affects adaptation to a non-stressful and a stressful environment in populations of the green alga Chlamydomonas reinhardtii, while minimizing the direct costs of sexual reproduction. Contrary to several previous studies, we found that an increasing frequency of sexual reproduction hindered adaptation of populations. In populations experiencing the highest frequency of sexual reproduction, adaptation was entirely prevented. These findings suggest that there were strong evolutionary costs associated with too frequent sexual reproduction in our populations. This observation may help to explain why in many facultative sexual species, there is a low frequency of sexual reproduction.

## Introduction

The geographical ranges that species occupy in the natural world are determined by how well those species are adapted to their abiotic environment (e.g. climatic conditions, soil composition) and their biotic environment (e.g. competitive interactions, predator-prey interactions, parasite-host interactions) (Kawecki and Ebert, 2004; Holt and Keitt, 2005; Bijlsma and Loeschcke, 2005; Hardie and Hutchings, 2010). One important mechanism that affects the potential to adapt is the reshuffling of genetic variation through sexual reproduction (Maynard-Smith, 1978; Bell, 1982). Sex can affect adaptation of species in several ways by for example purging deleterious mutations, aiding adaptation by bringing together and fixing novel adaptive mutations, or by recombining existing variation that can for example help in resisting parasite infections (Red Queen hypothesis)(reviewed in Hartfield and Keightley 2012).

Recent experimental work has demonstrated that sexual reproduction may speed up adaptation of species. For example, evolution experiments with populations of algae, protists, rotifers, and yeast have demonstrated that sexual reproduction can facilitate adaptation of populations to their biotic or abiotic environment (Colegrave, 2002; Goddard et al., 2005; Becks and Agrawal, 2012; Haafke et al., 2016b; McDonald et al., 2016; Luijckx et al., 2017; Petkovic and Colegrave, 2019; Moerman et al., 2020). Additionally, it has been shown experimentally that sexual reproduction is under positive selection when environmental complexity increases (Becks and Agrawal, 2013; Luijckx et al., 2017). Similarly, sexual reproduction has been shown to be advantageous in natural populations, for example by facilitating adaptation of species to the local environment by introgression of locally adapted genes (Currat et al., 2008), or by facilitating the escape from parasitism (Brockhurst et al., 2014). Despite these benefits of sexual reproduction, many species including plants (Dickinson et al., 2003), fungi (Nieuwenhuis and James, 2016), invertebrates (Suomalainen, 1950; Vershinina and Kuznetsova, 2016) and certain vertebrate species (Suomalainen, 1950; Neaves and Baumann, 2011) are capable of some form of facultative sexual reproduction or asexual reproduction. Moreover, in many of these species, sexual reproduction is only occasional. These observations suggests that while sex can facilitate adaptation, it also can be costly. These costs can come in two forms. Direct costs of sex are associated with the typically slower cell division during sexual reproduction in single-celled organisms, with the need for investment of resources in males (two-fold cost of sex), and the dilution of sexual reproduction alleles (cost of meiosis) (Maynard-Smith, 1978; Lively and Lloyd, 1990; Hartfield and Keightley, 2012; Williams, 2020). Evolutionary costs occur when sexual reproduction hinders adaptation because too frequent reshuffling of genetic material would break up adaptive allele combinations, and prevent effective selection. Whereas past theoretical work has investigated the direct costs extensively (though rarely experimentally, but see Gibson et al., 2017), the evolutionary costs of sexual reproduction have received less scrutiny (but see Becks and Agrawal, 2012). Theoretical work predicts that only occasional sexual reproduction is favourable over more frequent sexual reproduction (Green and Noakes, 1995; Hurst and Peck, 1996; Otto and Lenormand, 2002; Hadany and Comeron, 2008; Otto, 2009; D’Souza and Michiels, 2010), however experimental scrutiny of this prediction is currently largely lacking.

In this experiment, we investigated how the frequency of sexual reproduction affects adaptation of genetically diverse and outcrossed populations of the green alga *Chlamydomonas reinhardtii*. Specifically, we aimed to directly assess how the frequency of sexual reproduction affected evolutionary adaptation, under a situation where direct costs were minimised. To do so, we assessed how increasingly frequent sexual reproduction affected adaptation in a stressful environment (increased concentration of salt, previously used as a stressful environment (Lachapelle et al., 2015; Lachapelle and Bell, 2012); “salt lines”) and in a non stressful environment (non-elevated salt concentration; “no salt lines”), as well as correlated responses (i.e. presence of evolutionary trade-offs) between the different environments. We designed our experiment in such a way that the evolution lines experienced an approximately equal number of generations, independent of the frequency of sexual reproduction, to minimize direct costs associated with sexual reproduction. We then assessed how the frequency of sexual reproduction affected adaptation to the selection environment that populations experienced during experimental evolution.

### Hypotheses

Because we minimize direct costs associated with sexual reproduction, we could expect to see different relations between the frequency of sexual reproduction and the degree of adaptive evolutionary change. If sexual reproduction consistently facilitates adaptation, and there are no evolutionary costs associated with too frequent sexual reproduction, we would expect that the degree of adaptation increases directly with an increasing frequency of sexual reproduction (Figure 1, dark teal line). If, however, sexual reproduction is not costly, but too frequent sex would no longer aid adaptation, we would expect that the degree of adaptive evolutionary change initially increases quickly with the frequency of sexual reproduction, but this increase diminishes and levels off as the frequency of sexual reproduction increases further (Figure 1, light teal line). When too frequent sex would start to hinder adaptation, we expect that either the degree of adaptive evolutionary change starts to decrease if sexual reproduction would be too frequent (Figure 1, light brown line), or potentially even at low frequencies of sexual reproduction, entirely preventing any adaptive evolutionary change as sexual reproduction becomes more frequent (Figure 1, dark brown line). Based on theoretical predictions (Green and Noakes, 1995; Hurst and Peck, 1996; Otto and Lenormand, 2002; Hadany and Comeron, 2008; Otto, 2009; D’Souza and Michiels, 2010), we hypothesize that intermediate frequencies of sexual reproduction may have the most beneficial effect on adaptive evolutionary change, but too frequent sexual reproduction may become costly by breaking up beneficial allele combinations, and thus preventing effective selection (mild cost; Figure 1, light brown line). Because past experiments found that the advantages of sexual reproduction tend to be stronger in new or challenging environments (Colegrave, 2002; Kaltz and Bell, 2002; Goddard et al., 2005; Becks and Agrawal, 2012; Haafke et al., 2016b; McDonald et al., 2016; Luijckx et al., 2017; Petkovic and Colegrave, 2019; Moerman et al., 2020), we would however expect that in our experiment, sexual reproduction may be more advantageous for those populations that experienced evolution in the salt environment than in the no salt environment.

**Figure 1:**
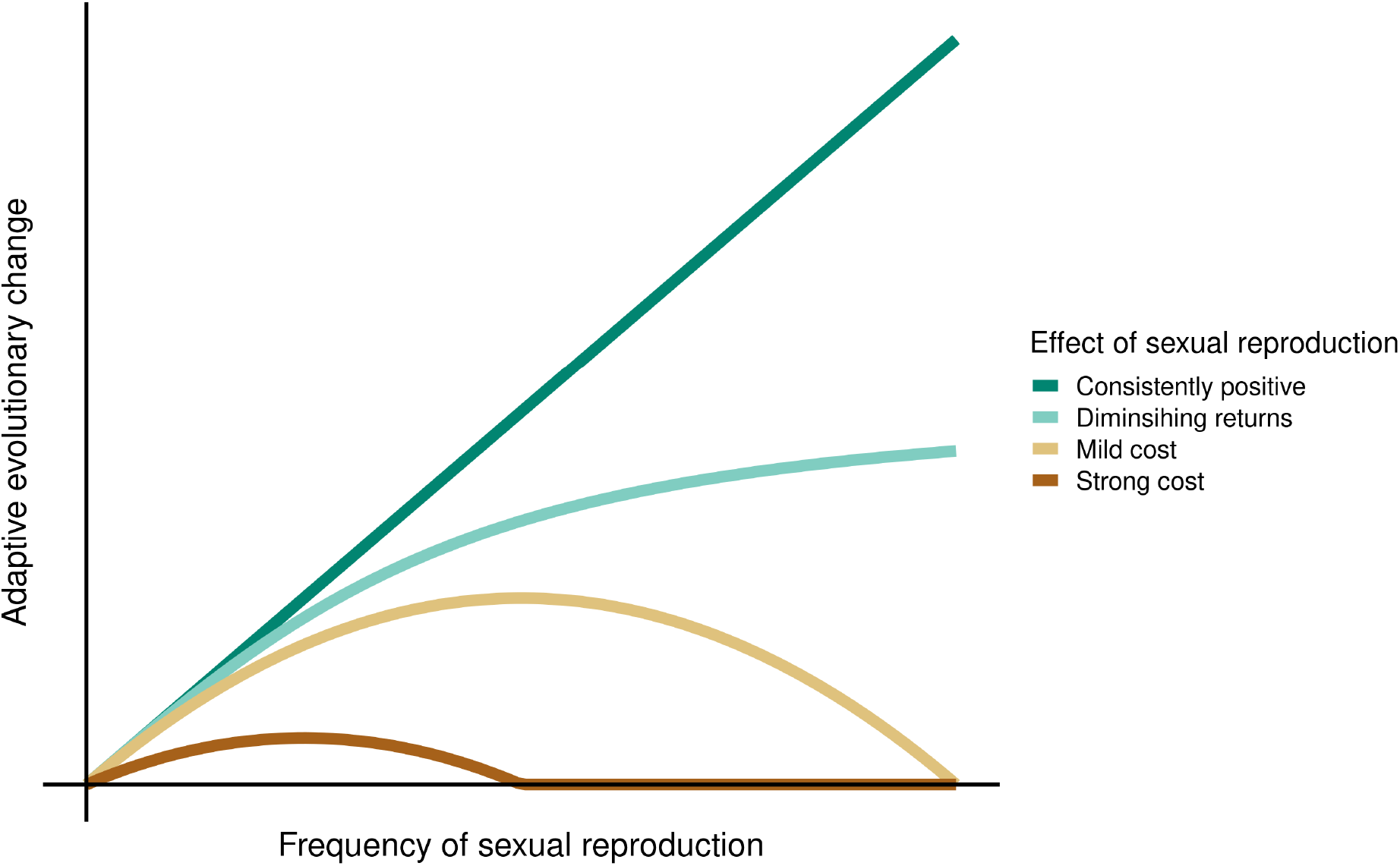
Hypothetical relation between the frequency of sexual reproduction and the degree of adaptive evolutionary change that a species may experience. We here show four hypothetical cases of a relation between the frequency of sexual reproduction and the degree of adaptive evolutionary change that we could expect to see in our experiment. Depending on the evolutionary costs and benefits, we could expect to see that either 1) sexual reproduction consistently facilitates adaptive evolutionary change (dark teal), 2) sexual reproduction has no evolutionary cost, but has diminishing return when species engage increasingly frequent in sexual reproduction (light teal), 3) too frequent sexual reproduction can have a mild cost, and reduces adaptive evolutionary change (light brown) or 4) where even low frequencies of sexual reproduction can become costly and start to reduce the degree of adaptive genetic change (dark brown). Note that this figure only shows evolutionary costs, not direct costs such as slower division rates.

**Figure 2:**
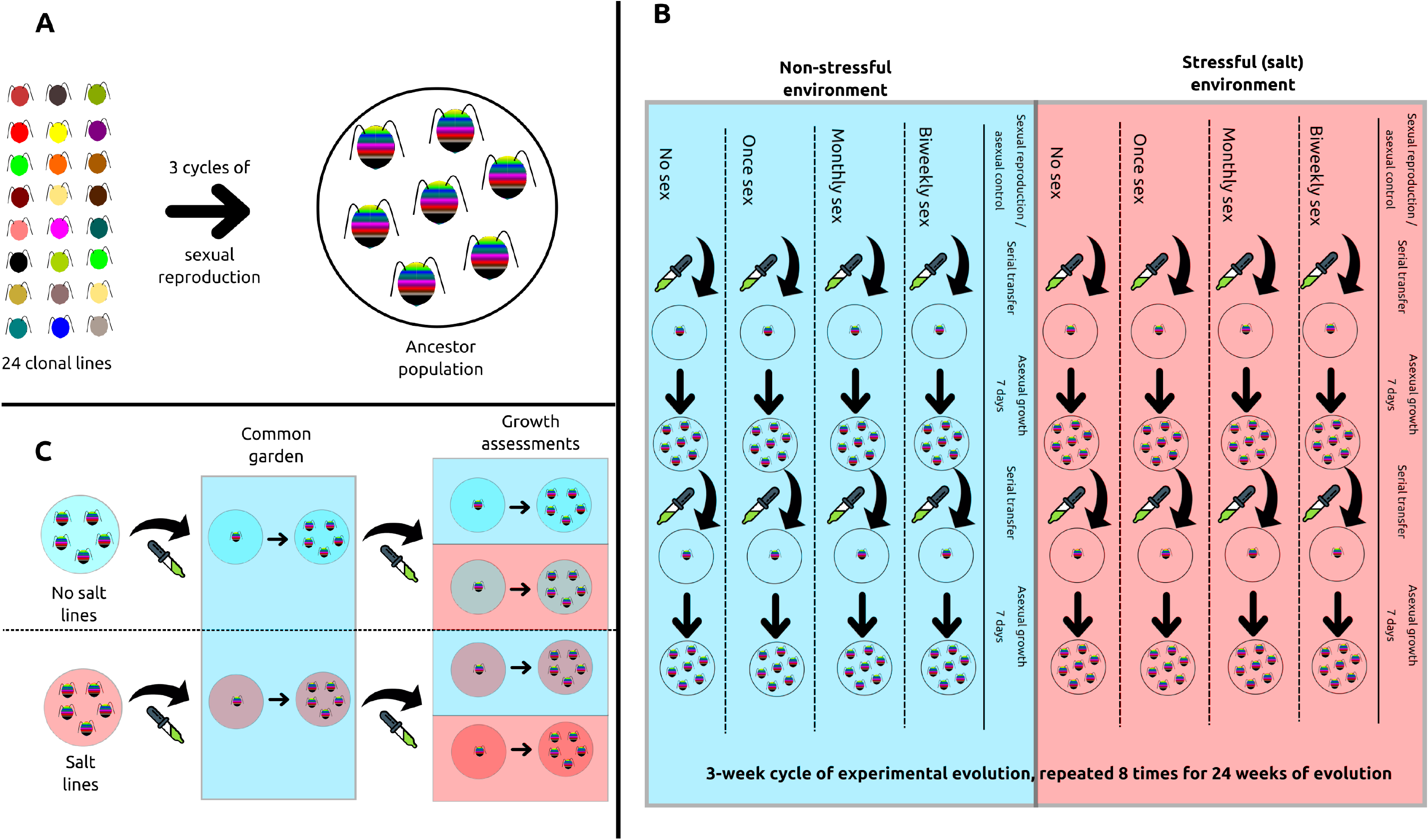
Schematic representation of the experimental setup. A) We created a genetically diverse and outcrossed ancestor population by mixing together 24 clonal lines of *Chlamydomonas reinhardtii*, and subjecting them to three cycles of sexual reproduction. B) We started the evolution experiment using this ancestor population, and subjected evolution lines either to a no salt environment (blue) or a salt environment (red). Evolution lines experienced different frequencies of sexual reproduction (none, once, monthly or biweekly). For each combination of the environment and frequency of sexual reproduction, we maintained six replicate evolution lines. The evolution lines experienced a total of 24 weeks of evolution, during which we repeated the same three week cycles, consisting of a sexual reproduction/asexual control phase, followed by two cycles of asexual growth. C) After experimental evolution, we subjected the evolution lines (blue circle=no salt lines; red circle=salt lines) to a common garden phase (no salt environment; blue box), and subsequently grew them in both environments (blue box=no salt environment; red box=salt environment) to measure adaptation to the selection environment.

## Materials and methods

### Model species and general culturing conditions

*Chlamydomonas reinhardtii* is a unicellular green alga, living in freshwater and soil environments. Because of its ease in culturing, short generation time, and strict control over its reproductive cycle, this species is commonly used in evolution experiments (Kassen and Bell, 1998; Collins and Bell, 2004; Ratcliff et al., 2013; Haafke et al., 2016a; Koch and Becks, 2017; Lachapelle and Colegrave, 2017; Böndel et al., 2019). We performed all experimental work with *C. reinhardtii* cultures under the same general conditions. We kept all cultures in a 23 °C incubator. During experimental evolution, we grew cultures either in 24 well plates, containing 2 mL of medium per well, or on agar plates containing 10 mL of Bold’s medium supplemented with 15 g L^−1^ of bacteriological agar (Harris, 2009, ; chapter 8). During fitness assays, we grew cultures in 96 well plates, containing 200 μL of medium per well. We kept 96 well plates and 24 well plates at all times on a shaker, rotating at 180 rpm.

### Ancestral population

We generated a genetically diverse and outcrossed ancestor population from 24 clonal strains of *C. reinhardtii* by subjecting them to three rounds of sexual reproduction (see the Supplementary Material Table S1 for a full list of all clones). To do so, we first grew all clones to equilibrium density in a 24 well plate. We followed an established protocol to induce mating of the *C. reinhardtii* cells (Lachapelle and Colegrave, 2017; Petkovic and Colegrave, 2019). We first mixed all 24 clones in a 50 mL Falcon tube, and centrifuged the Falcon tube for 10 minutes at 5000 rpm. We then decanted the supernatant, and resuspended cells in distilled water, to induce mating. Subsequently, we incubated the cells in a 24 well plate (2 mL of culture per well) until the next day. We checked whether cells were mating through the formation of a mating mat. We then transferred the mating mat in each of the wells using an inoculation loop to an agar plate containing Bold’s medium supplemented with 15 g L^−1^ of agar powder, and wrapped the plates in aluminium foil. We incubated the wrapped agar plates in the dark for four days. After this incubation period, we placed the agar plates in a −20 °C freezer, in order to kill non-mating cells. We subsequently removed the aluminium foil, and incubated the cells in the light for an additional two days. Following this incubation in the light, we added 5 mL of Bold’s medium to each of the agar plates, and left them for one hour to recover offspring cells from the mating. Next, we transferred 2 mL of medium from each agar plate to one well of a new 24 well plate, and incubated this plate for one week, in order for the populations to grow to equilibrium density. We then repeated this whole process (mixing all populations, incubation in nitrogen free medium, incubation in the dark, freezing and recovery of cells) two additional times to make sure populations were thoroughly outcrossed.

### Experimental evolution

In this evolution experiment, we aimed at assessing how the abiotic environment (non-stressful versus stressful environment) and the frequency of sexual reproduction affected adaptation of the ancestral population. Because we were mainly interested in the evolutionary costs and benefits of sexual reproduction, we aimed to minimize the direct costs associated with sexual reproduction. Because *Chlamydomonas* is an isogamous species, there is no cost of males present in this species. Additionally, the cost of meiosis has been argued to be rare and only present under very restrictive conditions (Treisman and Dawkins, 1976; Charlesworth, 1980), which do not apply to our model species. Therefore, the only direct cost of sexual reproduction in this system is the time needed for sexual reproduction, as the sexual reproductive cycle of *C. reinhardtii* takes much longer than asexual reproduction. Therefore we subjected populations that were not scheduled for sexual reproduction to an asexual control treatment (discussed below), aimed at ensuring that the number of generations was approximately similar for populations experiencing asexual or sexual reproduction.

### Experimental design and handling

We subjected a total of 48 replicate populations (from here on referred to as evolution lines) to experimental evolution. Half of those evolution lines experienced a no salt environment (Bold’s medium; hereafter call “no salt lines”), whereas the remaining half experienced evolution in a salt environment (Bold’s medium supplemented with 4 g L^−1^ NaCl; hereafter called “salt lines”). In each of the abiotic environments, we subjected the remaining 24 populations to four different frequencies of sexual reproduction: none (pure asexual reproduction), once (single sexual reproduction event, after 56 generations of asexual growth), monthly (sexual reproduction after every 4 weeks/28 generations of asexual growth) or biweekly (sexual reproduction after every 2 weeks/14 generations of asexual growth). We thus had six replicate evolution lines per treatment combination (see also Table S2 in the Supplementary Material). We subjected each of those evolution lines to a total of 24 weeks of experimental evolution, corresponding to approximately 112 generations of asexual growth, interspersed with eight sexual reproduction/asexual control events. These 24 weeks consisted of eight cycles of three weeks, during which the same steps were repeated in every cycle. Each cycle consisted of a first week during which the evolution lines experienced either sexual reproduction or an asexual control treatment. The remaining two weeks consisted each of an asexual growth cycle (asexual cell division). After these 24 weeks of experimental evolution, we subjected the evolution lines to a common garden treatment, after which we assessed the change in fitness of the evolution lines. Each of these handling steps is discussed in more detail below. See also Table S3 in the Supplementary Material for an overview of the handling in each week of the experiment.

### Sexual reproduction cycle and asexual control

To induce sexual reproduction, we transferred 2 mL of culture from the appropriate evolution lines (i.e. the evolution lines scheduled for sexual reproduction) to a 2 mL Eppendorf tube. We centrifuged those eppendorf tubes for 10 minutes at 5000 rpm in order to pellet the cells. We then decanted the supernatant, and resuspended cells in 2 mL of distilled water. Subsequently, we transferred the evolution lines to a new 24 well plate, which we incubated for one day. For evolution lines which were not scheduled for sexual reproduction (asexual control), we transferred 2 mL of culture directly to the new 24 well plate. Mating was visually confirmed through the formation of mating mats in the medium. After the 24 hours of incubation, we transferred the mating cells (mating mats) to an agar plate using an inoculation loop. For the asexual control, we instead pipetted 100 μL of culture directly on the agar plate. We subsequently wrapped the plates in aluminium foil, and incubated them for four days in the dark. During this incubation time, cells in the sexual reproduction treatment underwent meiosis, followed by a mitotic division. Due to the incubation in the dark, the rate of mitotic division of the evolution lines in the asexual control was strongly reduced. After this incubation period, we placed the agar plates for sexually reproducing populations in a −20 °C freezer for four hours in order to kill the asexual cells. We kept the agar plates with the evolution lines scheduled for asexual control in the incubator during this time. Afterwards, we removed the aluminium foil, and incubated the agar plates for an additional two days in the light. We then added 5 mL of medium to each of the agar plates (respectively Bold’s medium or Bold’s medium supplemented with 4 g L^−1^ NaCl) and left the plates to rest for one hour, in order to recover the cells. We then transferred 2 mL of culture to a new 24 well plate. Thus, sexual reproduction and asexual control only differed in the freezing and incubation in distilled water, but were otherwise identical, which should minimize differences in selection. This method resulted in approximately similar population densities between the evolution lines at the end of the sexual reproduction cycle/asexual control.

### Asexual growth cycle

To initiate an asexual growth cycle, we prepared fresh 24 well plates by adding medium to all the wells (2 mL of Bold’s medium for the non-stressful environment or 2 mL of Bold’s medium + 4 g L^−1^ NaCl for the stressful environment). We then transferred 20 μL of culture from the evolution lines to these new 24 well plates, and incubated these plates for one week (approximately seven generations). This asexual growth cycle was repeated twice (total of 14 days of asexual growth) in between every sexual reproduction/asexual control treatments.

### Common garden treatment

After experimental evolution, we subjected the evolution lines to a common garden environment, to reduce maternal and epigenetic effects. To do so, we transferred 20 μL of culture from the evolution lines to new 24 well plates containing Bold’s medium supplemented with 100 mg L^−1^ Ampicillin, to ensure all evolution lines were free from potential bacterial contamination. We chose the regular Bold’s medium for the common garden medium, as this was the same medium that the clonal populations were kept in prior to our evolution experiment. We subsequently incubated these common garden populations for one asexual growth phase (seven days). Thus, the evolution lines should have experienced a common garden environment for approximately 7 generations, prior to starting the population growth assays (see the section below).

### Population growth assays

To assess how the abiotic environment and the frequency of sexual reproduction experienced during evolution affected fitness change, we measured population growth of the evolution lines and the ancestor population in both abiotic environments (Bold’s medium or Bold’s medium + 4 g L^−1^ NaCl). For each of the evolved lines, we measured population growth of three replicate populations in each environment (total of 48 evolution lines × 2 environments × 3 replicates = 288 assays). For the ancestor population, we measured population growth of 36 replicate populations in each of both environment (2 environments × 36 replicates = 72 assays). We prepared population growth assays in 96 well plates, by adding 200 μL of medium to the wells, and inoculating the wells with 2 μL of culture from respectively the evolution lines or the ancestor population. To avoid drying out of the assays due to evaporation, we only used the central 60 wells of the 96 well plates for assays, and filled the wells of the outside rows and columns with medium only. Subsequently, we incubated the assays, and allowed them to grow for seven days, during which we measured population size twice per day (total of 14 absorbance measurements). Following established protocols (Lachapelle et al., 2015; Lachapelle and Colegrave, 2017; Petkovic and Colegrave, 2019), we measured optical density in the wells (OD_750_) as a proxy for population size. To account for background absorbance from the plates and medium, we subtracted for each plate the median absorbance of the empty wells (i.e. wells containing medium but no *Chlamydomonas* cells) from all absorbance measurements. For plots showing the standardized absorbance measurements, see section S5 in the Supplementary Material.

### Statistical analysis

We performed all statistical analyses using the R-statistical language version 4.1.2 (R Core Team, 2021).

#### Calculation of fitness change

In order to investigate fitness change of the evolution lines, we assessed two aspects of population growth: the intrinsic rate of increase (*r*_0_) and the maximum density that populations reached (equilibrium population density *K*). We chose these two metrics because the intrinsic rate of increase *r*_0_ has been found to be typically under selection in past experiments performed with this species, and because these metrics are independent of potential lag in growth that could occur during the assays. In order to estimate *r*_0_, we first calculated the growth rate between each two subsequent absorbance measurements *n*_1_ and *n*_2_ as:

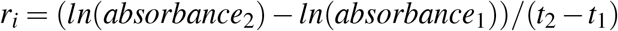

where *t*_1_ and *t*_2_ are the times since the start of the assays for the absorbance measurements. We then estimated *r*_0_ as the maximum value of all *r*_*i*_ values for each of the assays. Secondly, we calculated *K* as the maximum absorbance observed in each of the assays, over all 14 absorbance measurements.

In order to calculate change in fitness, relative to the ancestor population, we subsequently divided our *r*_0_ estimates and *K* estimates by the median value for the ancestor population. This allowed us to assess how the traits in the evolved lines had changed, relative to the ancestor, with a value of 1 indicating that evolution lines performed equally well as the ancestor populations, whereas positive (negative) values indicate an increase (decrease) in fitness.

#### Assessment of adaptation to the selective environment

To investigate how adaptation to the selective environment experienced during experimental evolution was affected by evolutionary history and the frequency of sexual reproduction, we assessed the change of fitness (intrinsic rate of increase and equilibrium population density) of evolved populations, in the assay environment that matched the environment they experienced during experimental evolution. That is, fitness change of no salt lines in Bold’s medium and fitness change of salt lines in Bold’s medium + 4 g L^−1^ NaCl. To do so, we first fit a linear mixed model (nlme package, version 3.1-155; Pinheiro et al., 2020), using evolutionary history (salt lines/no salt lines) and frequency of sexual reproduction (none/once/monthly/biweekly) as fixed effects, and population ID as a random effect. We subsequently ranked all possible models using the dredge function in the MuMIn package (version 1.43.17; Bartoń, 2009), based on the AICc criterion (Gelman et al., 2014). We selected the best fitting model, and report summary and type-III anova output. We do so separately for the intrinsic rate of increase (*r*_0_) and the maximum population density (*K*).

#### Assessment of correlated responses in both assay environments

To assess whether evolution lines experienced trade-offs in adaptation to the different environments, we next assessed the change of fitness (intrinsic rate of increase *r*_0_ and equilibrium population density *K*) of evolution lines, in both assay environments (salt environment and no salt environment). To do so, we fit a linear mixed model (nlme package, version 3.1-155; Pinheiro et al., 2020), using the assay environment (no salt environment/salt environment), evolutionary history (salt lines/no salt lines) and frequency of sexual reproduction (none/once/monthly/biweekly) as fixed effects, and population ID as a random effect. We then ranked all possible models based on the AICc criterion (Gelman et al., 2014) using the dredge function in the MuMIn package (version 1.43.17; Bartoń, 2009). Following model ranking, we selected the best fitting model (lowest AICc score), and report summary and type-III anova output of this best fitting model. We separately discuss the best fitting model for the intrinsic rate of increase (*r*_0_) and the maximum population density (*K*).

## Results

In this experiment, we investigated how the frequency of sexual reproduction affected adaptation of evolution lines subjected to approximately 100 generations of evolution in either a no salt environment or a salt environment. Specifically, we investigated how the intrinsic rate of increase *r*_0_ and the equilibrium population density *K* of evolution lines changed relative to the ancestor population, in the selection environment that populations experienced.

### Adaptation to the selection environment

#### Intrinsic rate of increase *r*_0_

The intrinsic rate of increase *r*_0_ was affected by the evolutionary history and the frequency of sexual reproduction, as well as the interaction between these two factors. For both the salt lines and the no salt lines, the proportional increase in *r*_0_, compared with the ancestor, declined with increasing frequency of sexual reproduction. In case of the no salt lines, (frequency of sexual reproduction; χ^2^_3_=19.172, p<0.001; Figure 3), the change in *r*_0_ was reduced by 0.11 and 0.35 for the evolution lines that experienced monthly or biweekly sexual reproduction, respectively. This negative effect of more frequent sexual reproduction was even more pronounced for salt lines (Evolutionary history × frequency of sexual reproduction; χ^2^_3_=8.041, p=0.045; Figure 3 right panel). Compared to the populations that experienced no sexual reproduction, the change in *r*_0_ of salt lines was reduced by respectively 0.12 if they experienced sexual reproduction once, by 0.38 if they experienced monthly sexual reproduction, and by 0.71 if they experienced biweekly sexual reproduction. Notably, both in the no salt lines and the salt lines, the evolution lines that experienced the highest frequency of sexual reproduction (biweekly) grew approximately as fast as the ancestor population, suggesting that adaptation was entirely prevented when experiencing a high frequency of sexual reproduction. Additionally, we observed that in the absence of sexual reproduction, *r*_0_ of salt lines increased more strongly than the *r*_0_ of no salt lines, relative to the ancestor (evolutionary history; χ^2^_1_=8.769, p=0.003; Figure 3). For full statistical output, see the Supplementary Material section S4.1.

**Figure 3:**
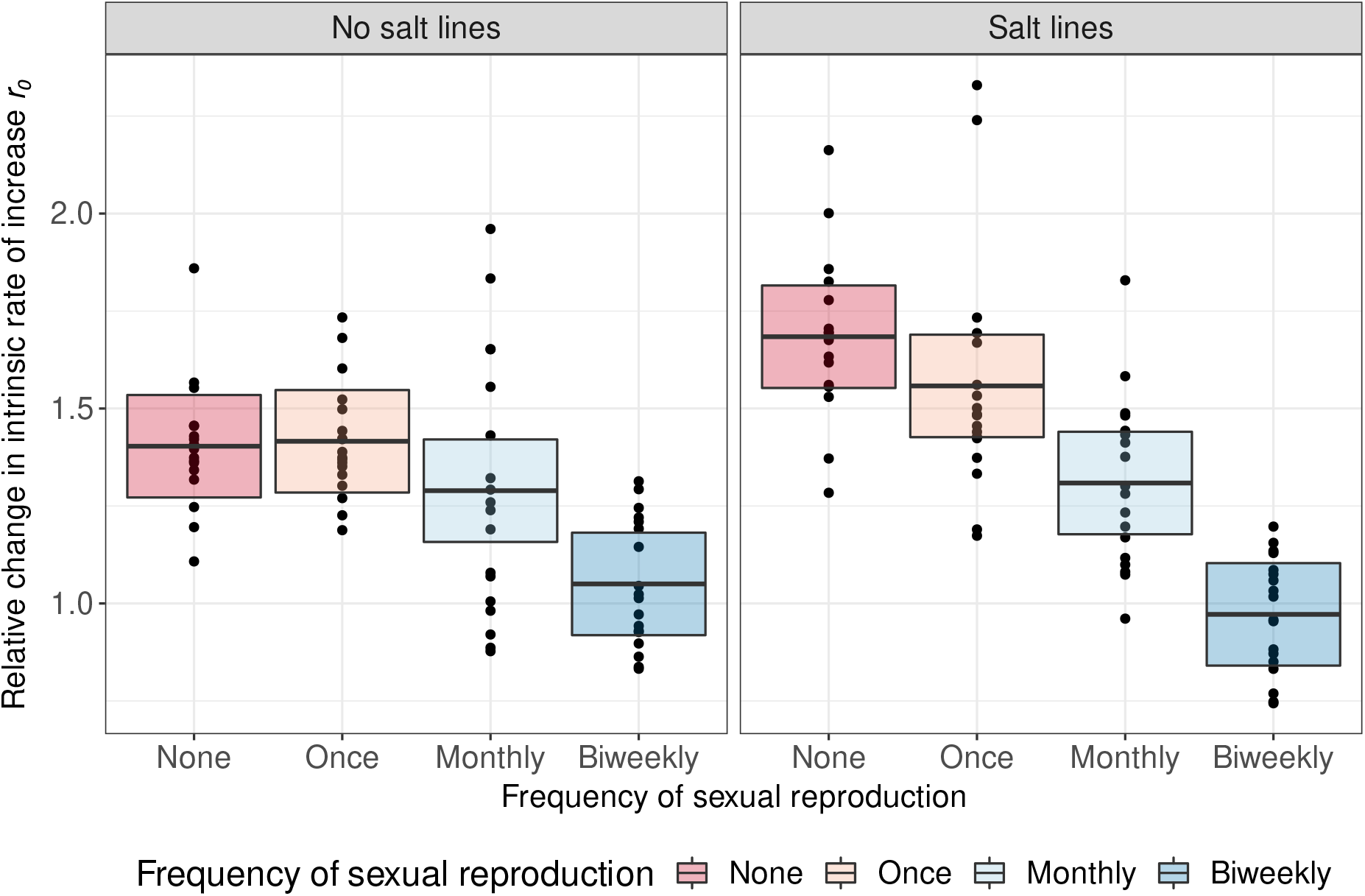
Higher frequency of sexual reproduction reduces change in the intrinsic rate of increase *r*_0_. The left panel shows data and model predictions for adaptation to the selection environment (intrinsic rate of increase) for the no salt lines, and the right panel for the salt lines. Circles represent individual measurements of change in intrinsic rate of increase of evolved lines, relative to the ancestor. Boxplots show the mean estimates (black lines) and 95 % confidence intervals (shaded areas) for the fixed effect estimates of the best fitting model. Colours represent the frequency of sexual reproduction during experimental evolution.

#### Equilibrium population density *K*

Contrary to the intrinsic rate of increase, we observed that the change in the equilibrium density *K* was not affected by the frequency of sexual reproduction. We observed however a clear effect of the evolutionary history, where salt lines showed stronger adaptation to the local environment in terms of the equilibrium population density than the no salt lines (Evolutionary history; χ^2^_1_=22.178, p<0.001; Figure 4). Full statistical output can be found in the Supplementary Material section S4.2.

**Figure 4:**
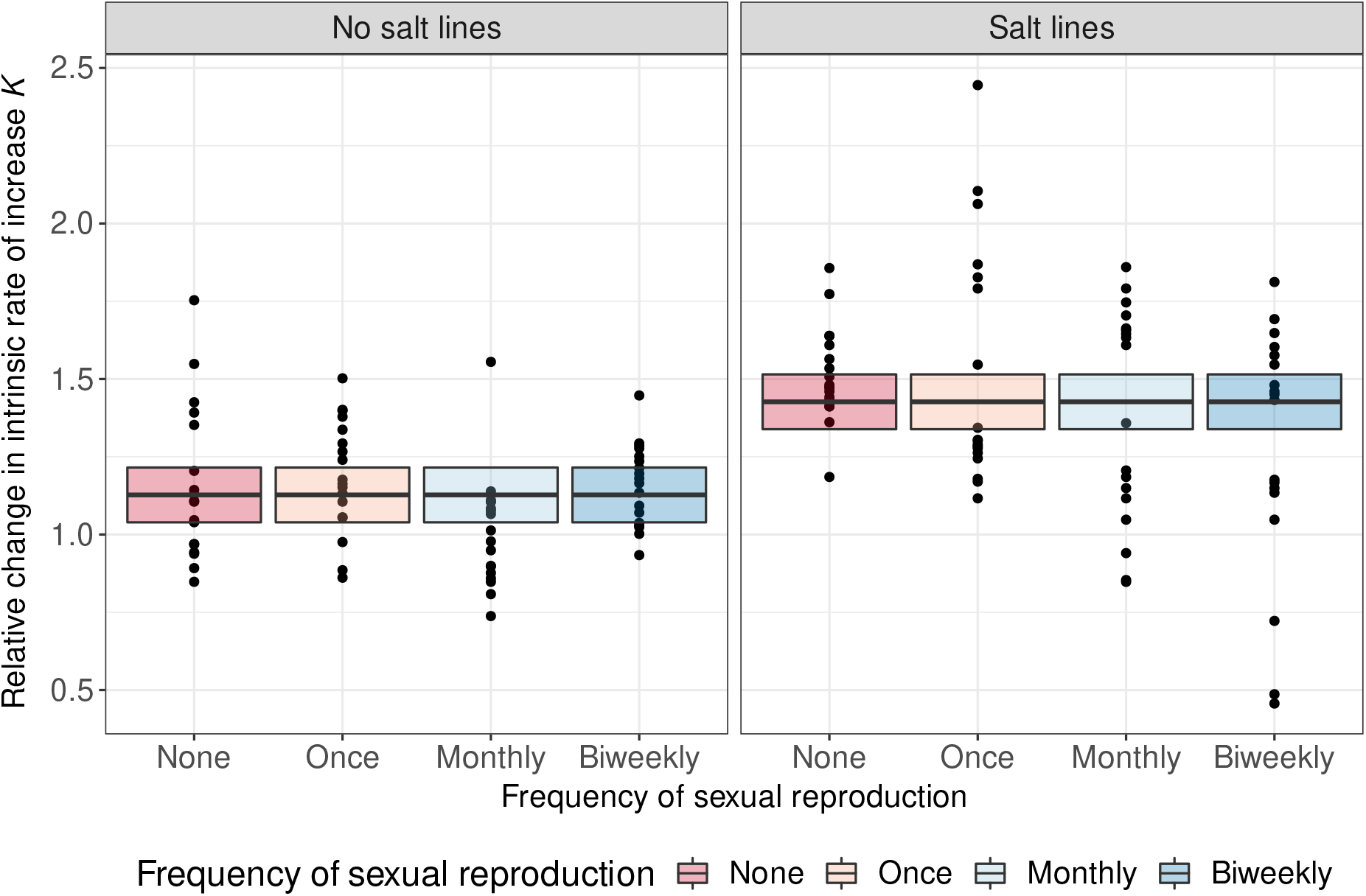
Evolutionary history shapes the change in the equilibrium density *K*. The left panel shows data and model predictions for adaptation to the local environment (equilibrium population density) for the no salt lines, and the right panel for the salt lines. Circles represent individual measurements of change in equilibrium population density of evolved lines, relative to the ancestor. Boxplots show the mean estimates (black lines) and 95 % confidence intervals (shaded areas) for the fixed effect estimates of the best fitting model. Colours represent the frequency of sexual reproduction during experimental evolution.

### Correlated responses in adaptation to the different selection environments

Next, to assess whether the evolution lines experienced any trade-offs in growth between the selection environment and the other environment, we investigated the correlated response of respectively the intrinsic rate of increase *r*_0_ and the equilibrium density *K* for both assay environments.

#### Intrinsic rate of increase *r*_0_

The change in the intrinsic rate of increase *r*_0_ was affected by the frequency of sexual reproduction, the evolutionary history of the evolution lines, as well as by the abiotic environment. More specific, we found that on average, the no salt lines increased more strongly in *r*_0_, irrespective of the abiotic environment (evolutionary history; χ^2^_1_=12.982, p=0.0003; Figure 5). Both salt lines and no salt lines grew significantly slower in the salt environment than in the no salt environment (abiotic environment; χ^2^_1_=21.631, p<0.0001; Figure 5). Independent of the evolutionary history and the abiotic environment, we found that an increasing frequency of sexual reproduction led to a smaller increase in *r*_0_ (frequency of sexual reproduction; χ^2^_3_=30.583, p<0.0001; Figure 5). We observed that the negative effect of frequent sexual reproduction was stronger for the salt lines than for the no salt lines (evolutionary history × frequency of sexual reproduction; χ^2^_3_=10.200, p=0.0169; Figure 5 right panels). However, we found no statistical indication of trade-offs in terms of the intrinsic rate of increase *r*_0_ (i.e. no significant interaction effect between the evolutionary history and the abiotic environment). Full statistical output can be found in section S4.3 of the Supplementary Material.

**Figure 5:**
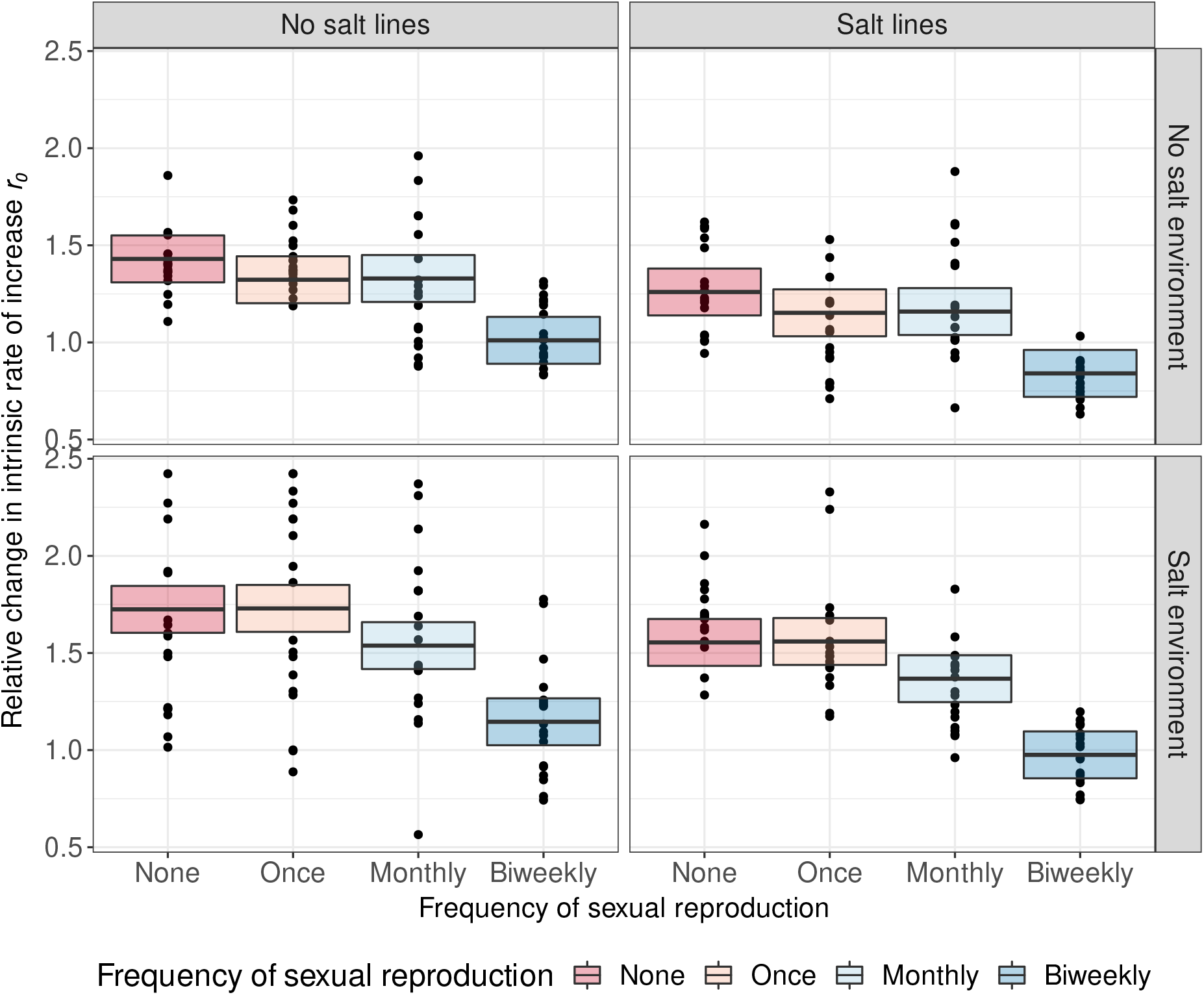
Higher frequency of sexual reproduction reduces change in the intrinsic rate of increase *r*_0_, independent of assay environment. The left panel shows data and model predictions for evolutionary trade-offs (intrinsic rate of increase) for the no salt lines, and the right panel for the salt lines. Circles represent individual measurements of change in intrinsic rate of increase of evolved lines, relative to the ancestor. Boxplots show the mean estimates (black lines) and 95 % confidence intervals (shaded areas) for the fixed effect estimates of the best fitting model. Colours represent the frequency of sexual reproduction during experimental evolution.

#### Equilibrium population density *K*

We observed that the equilibrium population density *K* was affected by the abiotic environment and the evolutionary history of the evolution lines, as well as their interaction. Specifically, we observed that *K* increased on average more strongly in the salt environment than in the no salt environment (abiotic environment; χ^2^_1_=11.202, p=0.0008; Figure 6). In the no salt environment, no salt lines showed a stronger increase in *K* than salt lines (evolutionary history; χ^2^_1_=4.741, p=0.030; Figure 6). However, in the salt environment, we observed the exact opposite patters, as the equilibrium population density *K* of salt lines increased more strongly than for the no salt lines (abiotic environment × evolutionary history; χ^2^_1_=23.213, p<0.0001; Figure 6). Overall, these results suggest that there is a trade-off in adaptation between the two environments, in terms of the equilibrium population density *K*. But we found no statistical indication that this trade-off was affected by the frequency of sexual reproduction in this data. See section S4.3 of the Supplementary Material for full statistical output.

**Figure 6:**
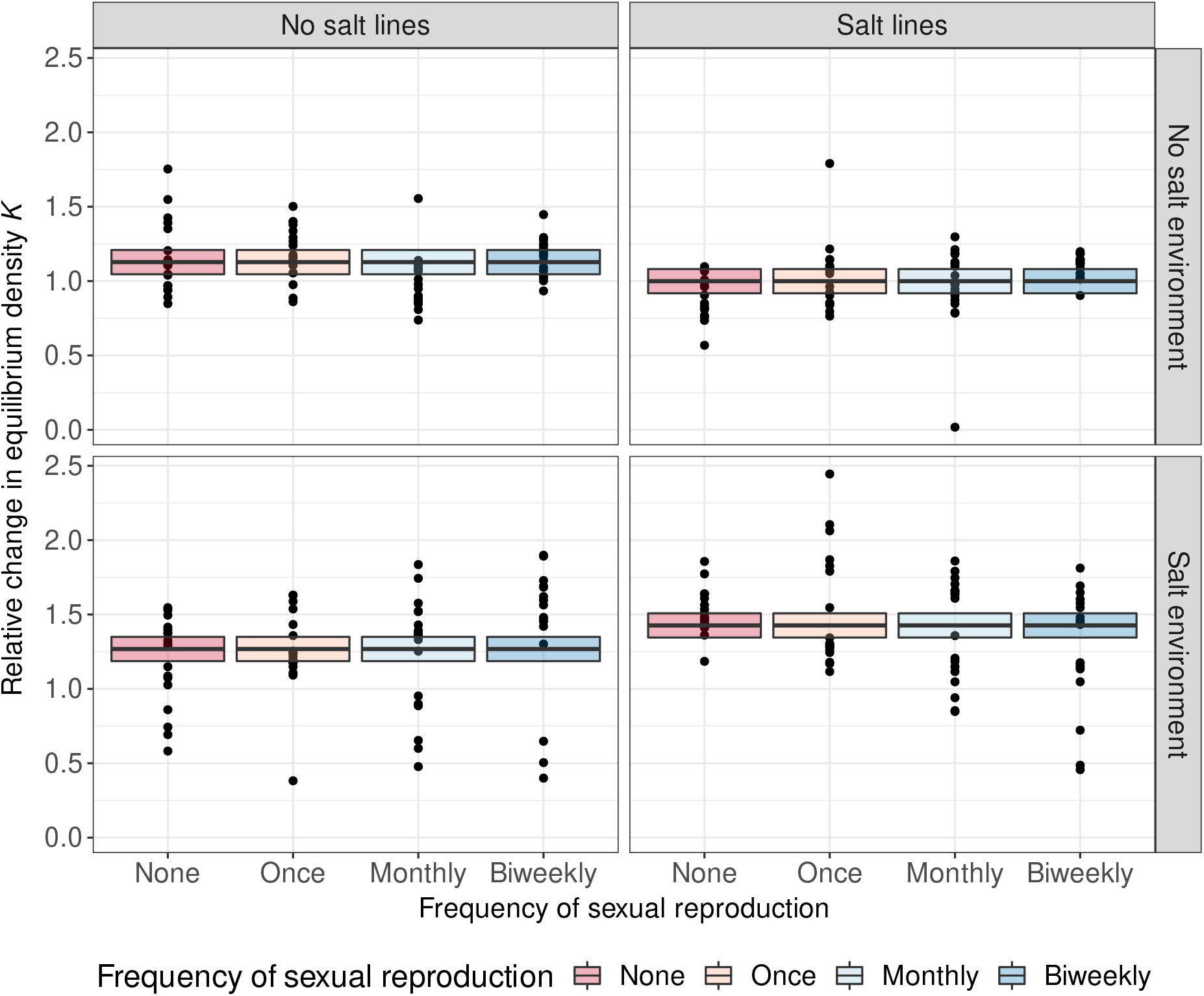
Evolutionary trade-offs affect the equilibrium density *K*. The left panels shows data and model predictions for the no salt lines, and the right panels for the salt lines. Top panels show data and model predictions in the no salt environment, whereas bottom panels show the salt environment. Circles represent individual measurements of change in equilibrium population density of evolved lines, relative to the ancestor. Boxplots show the mean estimates (black lines) and 95 % confidence intervals (shaded areas) for the fixed effect estimates of the best fitting model. Colours represent the frequency of sexual reproduction during experimental evolution.

## Discussion

We investigated the evolutionary costs and benefits of sexual reproduction in a no salt and a salt environment, while minimizing the direct costs associated with sexual reproduction. Specifically, we assessed how the frequency of sexual reproduction affected adaptation to the selection environment, as well as correlated responses in both assay environments (no salt environment and salt environment). We found that the frequency of sexual reproduction strongly affected adaptation to the selection environment in terms of the intrinsic rate of increase *r*_0_ in both the non stressful and the stressful environment (Figure 3). Specifically, an increasing frequency of sexual reproduction reduced adaptive evolutionary change in the evolution lines, up to the point where adaptation was entirely prevented in those populations that experienced the highest frequency of sexual reproduction. Surprisingly, we found that (in contrast to past experiments) sexual reproduction was not more advantageous in the challenging environment. Instead, sexual reproduction had a negative effect for lines that evolved in both the salt environment and the no salt environment. Contrary to the intrinsic rate of increase *r*_0_, adaptation to the selection environment in terms of the equilibrium population density *K* was not affected by the frequency of sexual reproduction. The lack of an effect of sexual reproduction on *K* may be because selection mostly acted on *r*_0_ and less on *K*, and therefore the effect of sexual reproduction was simply not significant. This explanation seems to be somewhat supported, as we saw little change in *K* for the no salt lines. For the salt lines, where there was a bit more change in *K*, it does appear as if there was less change in *K* when evolution lines experienced more frequent sexual reproduction (Figure 4, “Salt lines”), but the large variation between the replicates may have masked this effect statistically. When investigating correlated responses, we found that there were no trade-offs in terms of the intrinsic rate of increase *r*_0_. Rather, we found that a stronger increase in *r*_0_ in one assay environment corresponded to also a stronger increase in *r*_0_ in the second assay environment (Figure 5). This suggests that selection for growth mostly acted through a general growth response, rather than local adaptation to the selection environment. Contrary to the intrinsic rate of increase, we did find that evolution lines experienced a trade-off in terms of the equilibrium population density *K*. Specifically, we found that salt lines increased more in equilibrium population density in the salt assay environment than in the no salt assay environment, whereas the reverse was true for the no salt lines (Figure 6).

Our observation that an increasing frequency of sexual reproduction hinders adaptation during experimental evolution at first glance appears in contrast with our own prediction (see also Figure 1) and past theoretical work. Based on theoretical predictions (Green and Noakes, 1995; Hurst and Peck, 1996; Otto and Lenormand, 2002; Hadany and Comeron, 2008; Otto, 2009; D’Souza and Michiels, 2010), we would have expected to observe that a low to intermediate frequency of sexual reproduction is most beneficial. Additionally, we would have expected that higher frequencies of sexual reproduction would be more adaptive in those evolution lines that experienced a salt environment during experimental evolution. Previous experimental studies have found that sexual reproduction may speed up adaptation of populations, especially when they are subjected to complex or stressful environments (Colegrave, 2002; Kaltz and Bell, 2002; Goddard et al., 2005; Becks and Agrawal, 2012; McDonald et al., 2016; Luijckx et al., 2017; Petkovic and Colegrave, 2019; Moerman et al., 2020), but see Renaut et al. (2006). Whereas our observation may seem to be at odds with these previous studies, the cause of these differences may lie in the initial conditions of the experiment. In our current experiment, the ancestral population was both genetically diverse, and outcrossed. In this case, selection can likely act efficiently on this starting population, as the sexual reproduction prior to the start of the evolution experiment may have generated beneficial allele combinations from the mixed clonal lines. When such pre-adapted individuals would already have been present in the ancestral population, additional sexual reproduction may actually be detrimental. In this case, further sexual reproduction may break apart existing adaptive allele combinations, thus hindering adaptation (see for example Becks and Agrawal, 2012). This observation would also be in line with previous findings that up to three rounds of sexual reproduction may facilitate adaptation in *Chlamydomonas* populations, before sexual reproduction had diminishing returns on adaptation (Kaltz and Bell, 2002). Theoretical work has indicated that in such a case of well-mixed populations, sexual reproduction may be less advantageous, as it will no longer affect the genetic variation needed for effective selection (Otto, 2009). In contrast, several of the previous studies started out with populations which had either an extremely low genetic diversity (single or few clonal lines; Colegrave, 2002; Goddard et al., 2005; McDonald et al., 2016; Lachapelle and Colegrave, 2017) or with populations with a high degree of linkage disequilibrium (Moerman et al., 2020). Under these conditions, sexual reproduction may have played a more beneficial role. In case of the clonal populations, sexual reproduction may have played a beneficial role either by purging deleterious mutations or bringing together beneficial mutations/reducing clonal interference (Hill and Robertson, 1966; Gerrish and Lenski, 1998; Peck, 1994; Neiman et al., 2017). In the genetically more diverse populations but with a high degree of linkage disequilibrium, sexual reproduction may also have aided adaptation, by generating beneficial allele combinations from the existing genetic variation present in the different clonal populations (Otto and Barton, 1997; Burt, 2000; Otto and Lenormand, 2002). This is also in line with the theoretical prediction that sexual reproduction is mainly beneficial for populations by reducing selection interference between mutations/clonal lines (Hartfield and Keightley, 2012).

While our findings are in line with other theoretical and experimental studies, suggesting that too frequent sexual reproduction may hinder adaptation, it is still somewhat surprising that even relatively low frequencies of sexual reproduction could entirely halt any adaptation in our experimental populations. This observation may, however, help explain why facultative sexual species tend to engage only infrequently in sexual reproduction. Indeed, when looking at the natural world, many species have the capability to reproduce both asexually or sexually, and facultative sexual reproducing species typically engage in sexual reproduction only infrequently and when faced with adverse conditions (e.g. Hairston and Olds, 1984; Simon et al., 2002; Stelzer and Snell, 2003; Lynn and Doerder, 2012; Gerber et al., 2018). Whereas past studies have shown how sexual reproduction may be beneficial for adapting to new conditions, these studies for two reasons did not entirely elucidate why species would only engage infrequently in sexual reproduction. Firstly, although these studies show how sexual reproduction can aid adaptation, the starting populations from these experiments are often not representative of typical natural populations (due to the extremely low genetic diversity and strong linkage disequilibrium), and may be more similar to, for example, the conditions of invasions or small founder populations. Under such conditions, the benefits of sexual reproduction may be larger than in natural populations (e.g. Currat et al., 2008; Peischl and Excoffier, 2015). Secondly, given that these past results indicated that sexual reproduction strongly aided adaptation under those experimental conditions, they could not yet explain why there would be only infrequent sexual reproduction in natural populations. Such an intermediate to low frequency of sexual reproduction could either be caused by direct costs associated with sexual reproduction (e.g. slow cell division in single celled organisms, two-fold cost of sex, cost of meiosis Lively and Lloyd, 1990; Hartfield and Keightley, 2012; Williams, 2020), or due to evolutionary costs when sexual reproduction becomes too frequent. Whereas the direct costs are likely to play at least partially a role in reducing the frequency of sexual reproduction, they may be unlikely to entirely explain the observed frequency of sexual reproduction in facultative sexual species. Especially for populations that are near equilibrium density, and for which a slower reproduction is therefore likely less costly, the direct costs of sexual reproduction may be low, as also suggested by empirical observations (Suomalainen, 1950; Glesener and Tilman, 1978; Bell, 1982; Tilquin and Kokko, 2016; Williams, 2020) as well as some experimental work (Becks and Agrawal, 2013; Gibson et al., 2017). Thus, to explain the predicted and observed relatively low frequency of sexual reproduction of facultative sexually reproducing species, an additional explanation may be necessary in the form of an evolutionary cost due to too frequent sexual reproduction. Indeed as suggested by the results of our experiment, where we observed that sexual reproduction was hindering adaptation, even when we minimized the indirect costs of sexual reproduction, evolutionary costs due to too frequent sexual reproduction may play an important role in why many species only engage infrequently in sexual reproduction.

Although our experiment provides evidence that evolutionary costs due to too frequent sexual reproduction may limit adaptation, this in no way negates the existing compelling evidence from previous experimental studies that sexual reproduction may facilitate adaptation under certain conditions. As discussed above, the difference in these findings may stem from the initial conditions of these different evolution experiments. Consequently, there may exist a gradient of conditions in genetic diversity and the degree of linkage disequilibrium during which the effect of sexual reproduction shifts from beneficial for adaptation to hindering adaptation. Future work could further elucidate how the role of sexual reproduction hinges on these initial conditions. Furthermore, whereas the genetic composition of our experimental populations may be closer to natural populations, the experiment was nonetheless carried out in relatively simple experimental conditions (i.e. simple environmental stress). As such, these conditions may have been relatively easy for the populations to adapt to. Whereas several previous experiments found an adaptive effect of sex for populations adapting to similar environmental stressors (e.g., McDonald et al., 2016; Petkovic and Colegrave, 2019; Moerman et al., 2020), one experiment did indeed find that sexual reproduction does not become more common for genetically diverse populations adapting to a relatively simple environmental stressor (Luijckx et al., 2017), which is in line with our findings here that too frequent sex is not advantageous under those conditions. Sexual reproduction may become more beneficial when the environmental change is more complex, or when environmental conditions fluctuate over time. Future experiments could try to address how the benefits and costs of sexual reproduction change under these conditions. For example, using an experimental design that carefully controls the degree of genetic variation and degree of linkage disequilibrium, one can evaluate either how sexual reproduction alters adaptation, or how the frequency of sexual reproduction itself changes depending on these initial conditions. Such experiments could also assess the genetic diversity and linkage disequilibrium on a genetic level, something that was, due to practical constraints, not possible to do in this experiment. Additionally, evolution experiments could attempt to investigate how more realistic environmental conditions (e.g. multiple stressors, complex environmental stressors, temporally fluctuating environments) affect how sexual reproduction affects adaptation. Empirical studies may investigate natural populations of facultative sexually reproducing species, and try to assess whether the frequency of sexual reproduction is affected by the genetic compositions (standing genetic variation; linkage disequilibrium) of said populations, or the complexity of their environment. Additionally, future experimental work may further incorporate direct costs of sex into the equation (as already partially done by Becks and Agrawal (2013)), to see how this further alters the change in the frequency of sexual reproduction. In conclusion, we here demonstrated that too frequent sexual reproduction may lead to evolutionary costs that affect adaptation of populations to their environment, suggesting that the low frequency of sexual reproduction in natural populations may be, in part, due to such evolutionary costs. Future experimental endeavours may help in further elucidating the costs and benefits of sexual reproduction, thus advancing our understanding on when and why sex may be (dis)favoured in natural populations.

## Acknowledgements

F.M. was funded by the Swiss National Science Foundation, Early PostDoc Mobility Fellowship P2ZHP3_199658. We thank two anonymous reviewers for their helpful comments on a previous version of this manuscript.

