## Supplementary material for "Frequent sexual reproduction limits adaptation in outcrossed populations of the alga *Chlamydomonas reinhardtii*": SI

#### S1 Ancestor population

We generated a genetically outcrossed ancestor population by mixing together the clones listed below, and subjecting them to three rounds of sexual reproduction.

| | **Strain ID** | **Mating type** |
|---|---|---|
| 1 | CC-1373 | MT+ |
| 2 | CC-1690 | MT+ |
| 3 | CC-1952 | MT- |
| 4 | CC-2342 | MT- |
| 5 | CC-2344 | MT+ |
| 6 | CC-2931 | MT- |
| 7 | CC-2935 | MT- |
| 8 | CC-2936 | MT+ |
| 9 | CC-2937 | MT+ |
| 10 | CC-2938 | MT- |
| 11 | CC-3059 | MT+ |
| 12 | CC-3060 | MT+ |
| 13 | CC-3061 | MT- |
| 14 | CC-3062 | MT+ |
| 15 | CC-3065 | MT+ |
| 16 | CC-3067 | MT+ |
| 17 | CC-3068 | MT+ |
| 18 | CC-3069 | / |
| 19 | CC-3073 | MT- |
| 20 | CC-3084 | MT- |
| 21 | CC-3086 | MT+ |
| 22 | CC-3087 | MT- |
| 23 | CC-3089 | / |
| 20 | CC-3268 | MT- |

Table S1 Used *C. reinhardtii* clones to create the ancestral population. For each clone, we list its identifier (CC-number) and mating type, if known.

#### S2 Experimental treatments

| | **Abiotic environment** | **Frequency of sexual reproduction** | **Replicate ID's** |
|---|---|---|---|
| 1 | Bold's medium | None | 1-6 |
| 2 | Bold's medium | Once | 7-12 |
| 3 | Bold's medium | Monthly | 13-18 |
| 4 | Bold's medium | Biweekly | 19-24 |
| 5 | Bold's medium + 4 $g L^{-1}$ NaCl | None | 25-30 |
| 6 | Bold's medium + 4 $g L^{-1}$ NaCl | Once | 31-36 |
| 7 | Bold's medium + 4 $g L^{-1}$ NaCl | Monthly | 37-42 |

| | | | |
|---|---|---|---|
| 8 | Bold's medium + 4 $g L^{-1}$ NaCl | Biweekly | 43-48 |

Table S2 Summary of the experimental treatments of the evolution experiment. For each of the treatments, we list the abiotic environment that populations experienced during evolution, the frequency with which they experienced sexual reproduction, and the ID numbers for the replicate populations that underwent that specific treatment.

### S3 Handling during experimental evolution

| **Week** | **None** | **Once** | **Monthly** | **Biweekly** |
|---|---|---|---|---|
| **1** | Asexual control | Asexual control | Asexual control | Sexual reproduction |
| **2** | Asexual growth | Asexual growth | Asexual growth | Asexual growth |
| **3** | Asexual growth | Asexual growth | Asexual growth | Asexual growth |
| **4** | Asexual control | Asexual control | Sexual reproduction | Sexual reproduction |
| 5 | Asexual growth | Asexual growth | Asexual growth | Asexual growth |
| 6 | Asexual growth | Asexual growth | Asexual growth | Asexual growth |
| 7 | Asexual control | Asexual control | Asexual control | Sexual reproduction |
| 8 | Asexual growth | Asexual growth | Asexual growth | Asexual growth |
| 9 | Asexual growth | Asexual growth | Asexual growth | Asexual growth |
| 10 | Asexual control | Asexual control | Sexual reproduction | Sexual reproduction |
| 11 | Asexual growth | Asexual growth | Asexual growth | Asexual growth |
| 12 | Asexual growth | Asexual growth | Asexual growth | Asexual growth |
| 13 | Asexual control | Sexual reproduction | Asexual control | Sexual reproduction |
| 14 | Asexual growth | Asexual growth | Asexual growth | Asexual growth |
| 15 | Asexual growth | Asexual growth | Asexual growth | Asexual growth |
| 16 | Asexual control | Asexual control | Sexual reproduction | Sexual reproduction |
| 17 | Asexual growth | Asexual growth | Asexual growth | Asexual growth |
| 18 | Asexual growth | Asexual growth | Asexual growth | Asexual growth |
| 19 | Asexual control | Asexual control | Sexual reproduction | Sexual reproduction |
| 20 | Asexual growth | Asexual growth | Asexual growth | Asexual growth |
| 21 | Asexual growth | Asexual growth | Asexual growth | Asexual growth |
| 22 | Asexual control | Asexual control | Sexual reproduction | Sexual reproduction |
| 23 | Asexual growth | Asexual growth | Asexual growth | Asexual growth |
| 24 | Asexual growth | Asexual growth | Asexual growth | Asexual growth |
| 25 | Common garden | Common garden | Common garden | Common garden |
| 26 | Fitness assays | Fitness assays | Fitness assays | Fitness assays |

Table S3 Summary of the experimental handling related to the ”Frequency of sexual reproduction” treatments. Note that, except for the NaCl concentration in the medium, the handling procedure was the same for both abiotic environments (stressful and non stressful environment). We show in each week whether populations underwent asexual growth (regular cell division cycle in liquid medium), sexual reproduction, asexual control, common garden treatment or the fitness assays.

### S4 Statistical output

#### S4.1 Statistical output for the model investigating the effect of the evolutionary history and the frequency of sexual reproduction on adaptation to the local environment - the intrinsic rate of increase $r_0$.

| | **DF** | $\chi^2$**-value** | **Pr** ($>\chi^2$) |
|---|---|---|---|
| (Intercept) | 1 | 437.8898 | <0.0001 |
| Frequency of sexual reproduction | 3 | 19.1724 | 0.0002 |
| Evolutionary history | 1 | 8.7688 | 0.0031 |
| Frequency of sexual reproduction × Evolutionary history | 3 | 8.0409 | 0.0452 |

Table S4 Type III Anova table (fixed effects) for the best model describing adaptation to the local environment (the change of the intrinsic rate of increase ($r_0$)), according to AICc model comparisons. For each of the variables, we list the degrees of freedom (DF), the $\chi^2$-value, and the significance (Pr>$\chi^2$).

| | **Estimate** | **Std. Error** | **t-value** | **Pr (> $|t|$)** |
|---|---|---|---|---|
| (Intercept) | 1.4033 | 0.0671 | 20.9258 | <0.0001 |
| Frequency of sexual reproduction (once) | 0.0126 | 0.0948 | 0.1332 | 0.8947 |
| Frequency of sexual reproduction (monthly) | -0.1140 | 0.0948 | -1.2024 | 0.2363 |
| Frequency of sexual reproduction (biweekly) | -0.3531 | 0.0948 | -3.7236 | 0.0006 |
| Evolutionary history (Salt lines) | 0.2808 | 0.0948 | 2.9612 | 0.0051 |
| Frequency of sexual reproduction (once) × Evolutionary history (Salt lines) | -0.1388 | 0.1341 | 1.0349 | 0.3069 |
| Frequency of sexual reproduction (monthly) × Evolutionary history (Salt lines) | -0.2611 | 0.1341 | 1.9470 | 0.0586 |
| Frequency of sexual reproduction (biweekly) × Evolutionary history (Salt lines) | -0.3589 | 0.1341 | -2.6763 | 0.0107 |

Table S5 Summary table (fixed effects) for the best model describing adaptation to the local environment (the change of the intrinsic rate of increase ($r_0$)), according to AICc model comparisons. For each of the variables, we list the model estimate (Estimate), standard error on the estimate (Std. Error), the t-value and the significance (Pr (> $|t|$)).

##### S4.2 Statistical output for the model investigating the effect of the evolutionary history and the frequency of sexual reproduction on adaptation to the local environment - the equilibrium population density $K$.

| | **DF** | **$\chi^2$-value** | **Pr (>$\chi^2$)** |
|---|---|---|---|
| (Intercept) | 1 | 628.784 | <0.0001 |
| Evolutionary history | 1 | 22.178 | <0.0001 |

Table S6 Type III Anova table (fixed effects) for the best model describing adaptation to the local environment (the change of the equilibrium population density ($K$)), according to AICc model comparisons. For each of the variables, we list the degrees of freedom (DF), the $\chi^2$-value, and the significance (Pr>$\chi^2$).

| | **Estimate** | **Std. Error** | **t-value** | **Pr (> $\|t\|$)** |
|---|---|---|---|---|
| (Intercept) | 1.1273 | 0.0450 | 25.0756 | <0.0001 |
| Evolutionary history (Salt lines) | 0.2994 | 0.0636 | 4.7094 | <0.0001 |

Table S7 Summary table (fixed effects) for the best model describing adaptation to the local environment (the change of the equilibrium population density ($K$)), according to AICc model comparisons. For each of the variables, we list the model estimate (Estimate), standard error on the estimate (Std. Error), the t-value and the significance (Pr (> $|t|$)).

##### S4.3 Statistical output for the model investigating the effect of the evolutionary history and the frequency of sexual reproduction on evolutionary trade-offs - the intrinsic rate of increase $r_0$.

| | **DF** | **$\chi^2$-value** | **Pr (>$\chi^2$)** |
|---|---|---|---|
| (Intercept) | 1 | 538.567 | <0.0001 |
| Frequency of sexual reproduction | 3 | 30.583 | <0.0001 |
| Evolutionary history | 1 | 12.982 | 0.0003 |
| Abiotic environment | 1 | 21.631 | <0.0001 |
| Abiotic environment × Frequency of sexual reproduction | 3 | 10.200 | 0.0169 |

Table S8 Type III Anova table (fixed effects) for the best model describing trade-offs between adaptation to the different environments (the change of the intrinsic rate of increase ($r_0$)), according to AICc model comparisons. For each of the variables, we list the degrees of freedom (DF), the $\chi^2$-value, and the significance (Pr>$\chi^2$).

| | **Estimate** | **Std. Error** | **t-value** | **Pr ($>\|t\|$)** |
|---|---|---|---|---|
| (Intercept) | 1.4299 | 0.0616 | 23.2070 | <0.0001 |
| Evolutionary history (Salt lines) | -0.1703 | 0.0473 | -3.6030 | 0.0008 |
| Abiotic environment (Salt environment) | 0.2948 | 0.0634 | 4.6509 | <0.0001 |
| Frequency of sexual reproduction (Once) | -0.1073 | 0.0805 | -1.3327 | 0.1896 |
| Frequency of sexual reproduction (Monthly) | -0.1008 | 0.0805 | -1.252 | 0.2173 |
| Frequency of sexual reproduction (Biweekly) | -0.4191 | 0.0805 | -5.2090 | <0.0001 |
| Evolutionary history (Salt lines) × Frequency of sexual reproduction (Once) | 0.1120 | 0.0896 | 1.2503 | 0.2124 |
| Evolutionary history (Salt lines) × Frequency of sexual reproduction (Monthly) | -0.0859 | 0.0896 | -0.9580 | 0.3391 |
| Evolutionary history (Salt lines) × Frequency of sexual reproduction (Biweekly) | -0.1597 | 0.0896 | -1.7814 | 0.0761 |

Table S9 Summary table (fixed effects) for the best model describing trade-offs between adaptation to the different environments (the change of the intrinsic rate of increase ($r_0$)), according to AICc model comparisons. For each of the variables, we list the model estimate (Estimate), standard error on the estimate (Std. Error), the t-value and the significance (Pr ($>|t|$)).

##### S4.4 Statistical output for the model investigating the effect of the evolutionary history and the frequency of sexual reproduction on evolutionary trade-offs - the equilibrium population density $K$.

| | **DF** | $\chi^2$**-value** | **Pr** ($>\chi^2$) |
|---|---|---|---|
| (Intercept) | 1 | 735.4554 | <0.0001 |
| Evolutionary history | 1 | 4.7406 | 0.0295 |
| Abiotic environment | 1 | 11.2020 | 0.0008 |
| Abiotic environment × Evolutionary history | 1 | 23.2125 | <0.0001 |

Table S10 Type III Anova table (fixed effects) for the best model describing trade-offs between adaptation to the different environments (the change of the equilibrium population density ($K$)), according to AICc model comparisons. For each of the variables, we list the degrees of freedom (DF), the $\chi^2$-value, and the significance (Pr>$\chi^2$).

| | **Estimate** | **Std. Error** | **t-value** | **Pr (> $\|t\|$)** |
|---|---|---|---|---|
| (Intercept) | 1.1273 | 0.0416 | 27.1193 | <0.0001 |
| Evolutionary history (Salt lines) | -0.1280 | 0.0588 | -2.1772 | 0.0346 |
| Abiotic environment (Salt environment) | 0.1408 | 0.0421 | 3.3469 | 0.0009 |
| Evolutionary history (Salt lines) × Abiotic environment (Salt environment) | 0.2866 | 0.0595 | 4.8179 | <0.0001 |

Table S11 Summary table (fixed effects) for the best model describing trade-offs between adaptation to the different environments (the change of the equilibrium population density ($K$)), according to AICc model comparisons. For each of the variables, we list the model estimate (Estimate), standard error on the estimate (Std. Error), the t-value and the significance (Pr ($>|t|$)).

#### S5 Optical density measurements during population growth assays

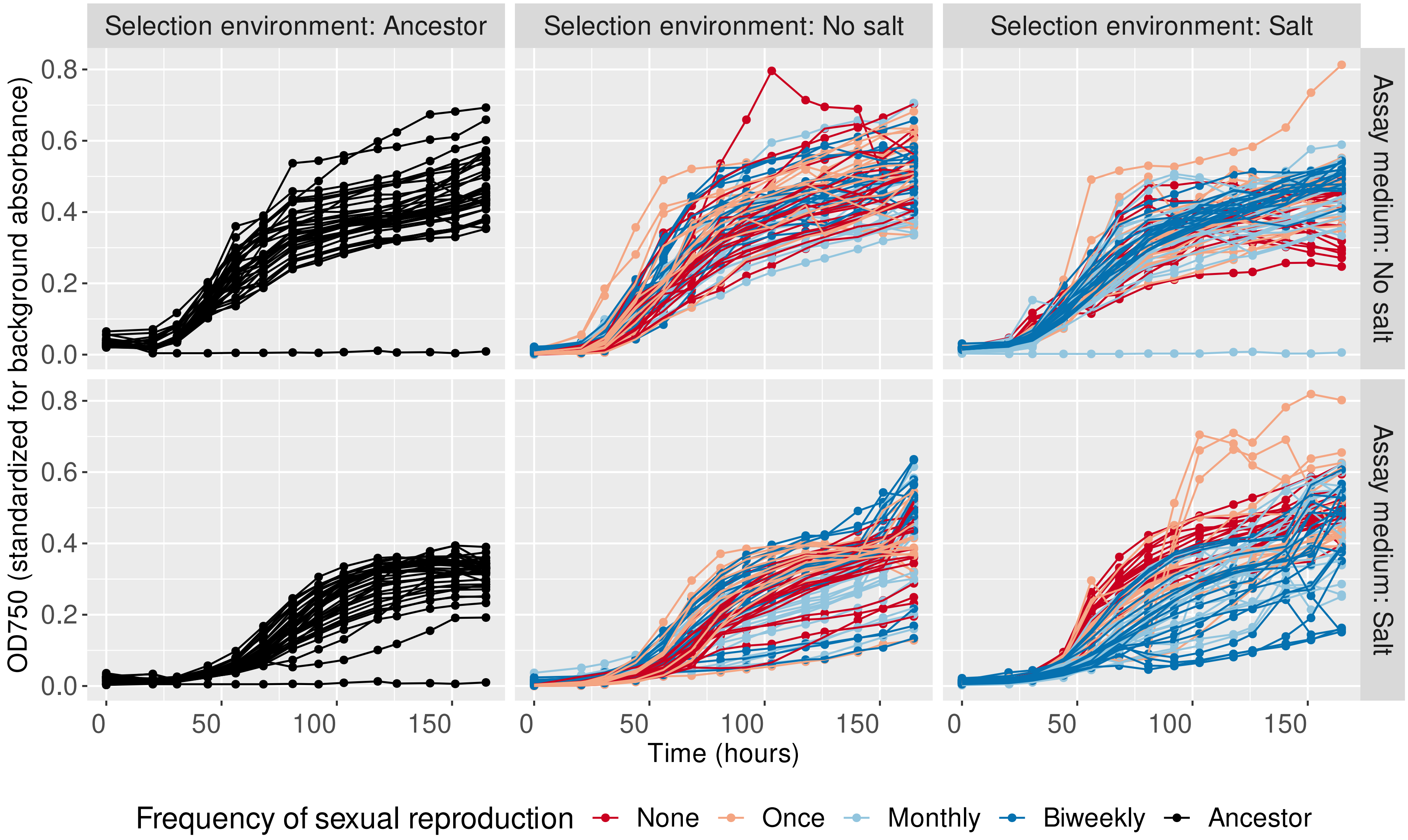


Figure S1: **Standardized optical density measurements for the population growth assays.** The plots show the optical density (absorbance at 750 nm), standaridzed for the background absorbance from the medium without cells (y-axis) over time (x-axis). Each line represents a single assay. The subplots show measurments for the ancestor population (left), the evolution lines that experienced evolution in the medium without added salt (middle plots; no salt lines) and the evolution lines that experienced evolution in medium without added salt (right plots; no salt lines).
